# A Secreted Subunit SARS-CoV-2 RBD-CpE Yeast Oral Vaccine Induces an Adaptive Immune Response in BALB/c Mice

**DOI:** 10.64898/2025.12.08.692901

**Authors:** John Donnie A. Ramos, Christopher Bresnahan, Taylor Brysgel, Noah Kozub, Gabriel Mendoza, Lexi Caitlin Murillo, Mark Kevin Devanadera, Josefino R. Castillo, Daniel Albert E. Castillo, Jose Paolo Aguilar, Marilyn G. Rimando, Nicanor Austriaco

**Affiliations:** Department of Biological Sciences, College of Science, University of Santo Tomas, Manila, PHILIPPINES; Department of Biology, Providence College, Providence, Rhode Island 02918, USA; Research Center for the Natural and Applied Sciences, University of Santo Tomas, Manila, PHILIPPINES; UST Laboratory for Vaccine Science, Molecular Biology, and Biotechnology, University of Santo Tomas, Manila, PHILIPPINES; Department of Biochemistry, Faculty of Pharmacy, University of Santo Tomas, Manila, PHILIPPINES

**Keywords:** yeast oral vaccine, COVID-19, oral vaccine, SARS-CoV2, Saccharomyces boulardii Introduction

## Abstract

Despite their differences, all the current COVID-19 vaccines need to be refrigerated and administered intramuscularly by a health care worker. They are also relatively expensive. These characteristics often make COVID-19 vaccine storage, distribution, and administration relatively complex, especially in low- and middle-income countries where refrigeration and healthcare workers are limited. To address these challenges, we are developing a COVID-19 oral vaccine utilizing the yeast Saccharomyces boulardii as a delivery platform. We have successfully engineered recombinant strains of Saccharomyces boulardii to express and secrete the Receptor Binding Domain (RBD) of the Spike protein from the original Wuhan-Hu-1 strain of the SARS-CoV-2 virus, fused to the C-terminal fragment of Clostridium perfringens enterotoxin (CpE). Animal trials suggest that our candidate secreted oral yeast vaccine can elicit detectable RBD-specific IgG and IgA, as well as an IFN-γ but not an IL-4 response, in BALB/c mice. As such, our data suggest that Saccharomyces boulardii remains a promising and novel vaccine delivery platform not only for COVID-19 but also for other infectious diseases.

## Introduction

According to the World Health Organization, as of November 1, 2025, a total of 13.64 billion COVID-19 vaccine doses have been administered worldwide (https://covid19.who.int/). These COVID-19 vaccines have been developed with both classical and novel vaccine platforms (1–6). Despite their differences, however, all the current COVID-19 vaccines require refrigeration and must be administered intramuscularly by a healthcare worker. They are also relatively expensive. These characteristics often make COVID-19 vaccine storage, distribution, and administration relatively challenging, especially in low- and middle-income countries where refrigeration and healthcare workers are limited (7).

To address these challenges, we are developing a COVID-19 oral vaccine utilizing the yeast Saccharomyces boulardii as a delivery platform. Yeast oral vaccines are being developed against a diverse range of diseases, including both animal and human diseases, as well as COVID-19 (8). Vaccines that are delivered orally have several advantages over their counterparts that are administered via injection (9, 10). Physiologically, the oral delivery of vaccines can stimulate both humoral and cellular immune responses at both systemic and mucosal sites, which, in principle, should establish broader and longer-lasting protection (11). Oral vaccines are also non-invasive, convenient, and can be self-administered. These are characteristics of a vaccine that can help increase immunization coverage in a population during a pandemic. It is striking that a recent study of over 15,000 adults in the United Kingdom revealed that fears of injection can explain 10% of COVID-19 vaccine hesitancy (12).

In this paper, we report that we have successfully engineered recombinant strains of Saccharomyces boulardii to express and to secrete the Receptor Binding Domain (RBD) of the Spike protein of the original Wuhan-Hu-1 strain of the SARS-CoV2 virus, fused to the C-terminal fragment of the Clostridium perfringens enterotoxin (CpE) that targets the tight junctions of mucosal associated lymphoid tissues (13–15). Animal trials suggest that our candidate vaccine can elicit detectable RBD-specific IgG and IgA, as well as IFN-γ but not an IL-4 response, in BALB/c mice. As such, our data suggest that Saccharomyces boulardii remains a promising and novel vaccine delivery platform not only for COVID-19 but also for other infectious diseases.

## Materials and Methods

### Yeast Strains and Growth Conditions

All experiments were done with the M2 Saccharomyces boulardii strain (16). Cells were cultured and treated using standard yeast media and protocols, as detailed in the Cold Spring Harbor yeast handbook (17). Yeast cells used for oral gavage of mice were grown in YPD. Unless noted otherwise, all drugs and reagents were purchased from SIGMA-Aldrich.

### Plasmid Design

The plasmid, pYIP-URA3-TEF1>{MF-6HIS-SARS-CoV2(RBD)-G4S-CPE}, which was used to express and to secrete the SARS-CoV2 RBD of the Spike protein in yeast was constructed and verified by VectorBuilder.com (Vector ID: VB201120-1148vcu). The expression cassette contains the following elements: TEF1, the Saccharomyces cerevisiae translation elongation factor EF-1α promoter; MF-6HIS-SARS-CoV2(RBD)-G4S-CPE, an ORF for the 6-His tagged receptor binding domain of the SARS-CoV-2 Wuhan-Hu-1 strain, with the yeast mating factor secretion signal at its N-terminus, and with the claudin-targeting sequence from Clostridium perfringens enterotoxin (CpE) at its C-terminus; and the CYC1 transcription terminator. A map of the plasmid, along with its sequence, can be obtained from the VectorBuilder.com website.

### Yeast Transformation

The pYIP-URA3-TEF1>{MF-6HIS-RBD(SarsCoV2)-G4S-CPE} plasmid was linearized with StuI and transformed into the M2 Saccharomyces boulardii strain with the standard LiAc/SS carrier DNA/PEG method (18) with the following modification: After the initial 30 minutes of incubation at 30°C, 40μL of DMSO was added to each transformation mix. The yeast cells were then heat-shocked at 42°C for 15 minutes, before being spun down and washed in sterile, deionized water. Finally, the cells were resuspended in 1mL of synthetic defined (SD) medium, 200μL of which was plated on SD plates lacking uracil to select for transformants.

### Collection and Concentration of Secreted RBD-CPE Antigen from Spent Media

Single colonies of transformed S. boulardii were cultured in 100mL of synthetic defined (SD) media for 48 hours at 37°C. After 15mL of the cultured cells were spun down, the supernatant/spent media was collected and concentrated using 50mL Amicon Ultra-15 Centrifugal Filter Units with a 30 kDa MWCO (Millipore), spun at 3550 RCF at 4°C for 30 minutes in a Centrifuge 5810R (Eppendorf).

### Western Blot Analysis

15μL of the protein concentrate obtained from spent yeast media was denatured in 2- mercaptoethanol/4x Laemmli loading sample (Bio-Rad) by incubating each sample at 95°C for 5 minutes. Samples were then run on a 10% Mini-PROTEAN TGX Precast Protein Gel (Bio-Rad) using a 10% SDS running buffer for 60 minutes at 150V (Mini Trans-Blot Cell from Bio-Rad). 15μL of a broad-spectrum color standard (New England Biolabs) was loaded as a ladder. SARS-CoV-2 (2019-nCoV) Spike RBD-His Recombinant Protein (SinoBiological) was used as a positive control at a 1:50 dilution. The gel was then wet transferred for 60 minutes onto a nitrocellulose membrane at 100 V/350 mA using a 20% methanol solution in 1X Tris/Glycine buffer in a Mini Trans-Blot Cell from Bio-Rad. The membrane was dried at 37°C and blocked for one hour in TBS- T 5% milk at room temperature on a slow rocking platform. The membrane was rinsed three times for 15 minutes in TBS-T and probed overnight with SARS-CoV-2 Spike Protein (RBD) Polyclonal Antibody (ThermoFisher) at a 0.35 μL/mL dilution in TBS-T with 1% milk at 4°C. After another three rinses in TBS-T, the membrane was incubated at room temperature with goat anti-rabbit IgG (H+L) Cross-Adsorbed Secondary Antibody, Alexa Fluor 700 (ThermoFisher) in Intercept T20 (TBS) Antibody Diluent (LI-Cor) at a 1:1000 dilution for one hour. The membrane was washed three times for 15 minutes in TBS-T and imaged with a LI-COR Odyssey DLx.

### Animal Trials

Twenty-four BALB/c mice were purchased from MOTS Animal House (Laguna, Philippines) and were divided into four groups of six mice each. They were housed at the College of Science animal facility at the University of Santo Tomas. The first two animal trials included cohorts of BALB/c mice inoculated with *S. boulardii* M2 alone, with *S. boulardii* M2 expressing GFP, or with one of two candidate vaccine S. boulardii strains (W2 or W5) that expressed and secreted RBD-CpE. Given the results of the first two trials, which suggested that both recombinant yeasts were comparable, and that both negative controls were comparable, we decided to limit our third and fourth animal trials to the W5-RBD vaccine mice alone, with a wild-type M2 yeast with no insert alone as a negative control.

As shown in Figure 1, on days 1-3, the cohorts of mice received their designated strain of S. boulardii as a primary dose by oral gavage using a #8 gavage needle. An oral dose of 2x109 yeast cells/mouse resuspended in 1mL PBS was used (19). Oral administration of the yeast was repeated on days 13-15 as a booster dose. Blood samples and feces were collected from all mice on days 0, 13, and 25. At the end of the fourth animal trial, spleens were also harvested on Day 25. This oral immunization protocol is similar to published protocols in the scientific literature (19, 20). Our animal study was approved by the UST-IACUC on September 28, 2022, with protocol number RC2021-030723.

**Figure 1.**
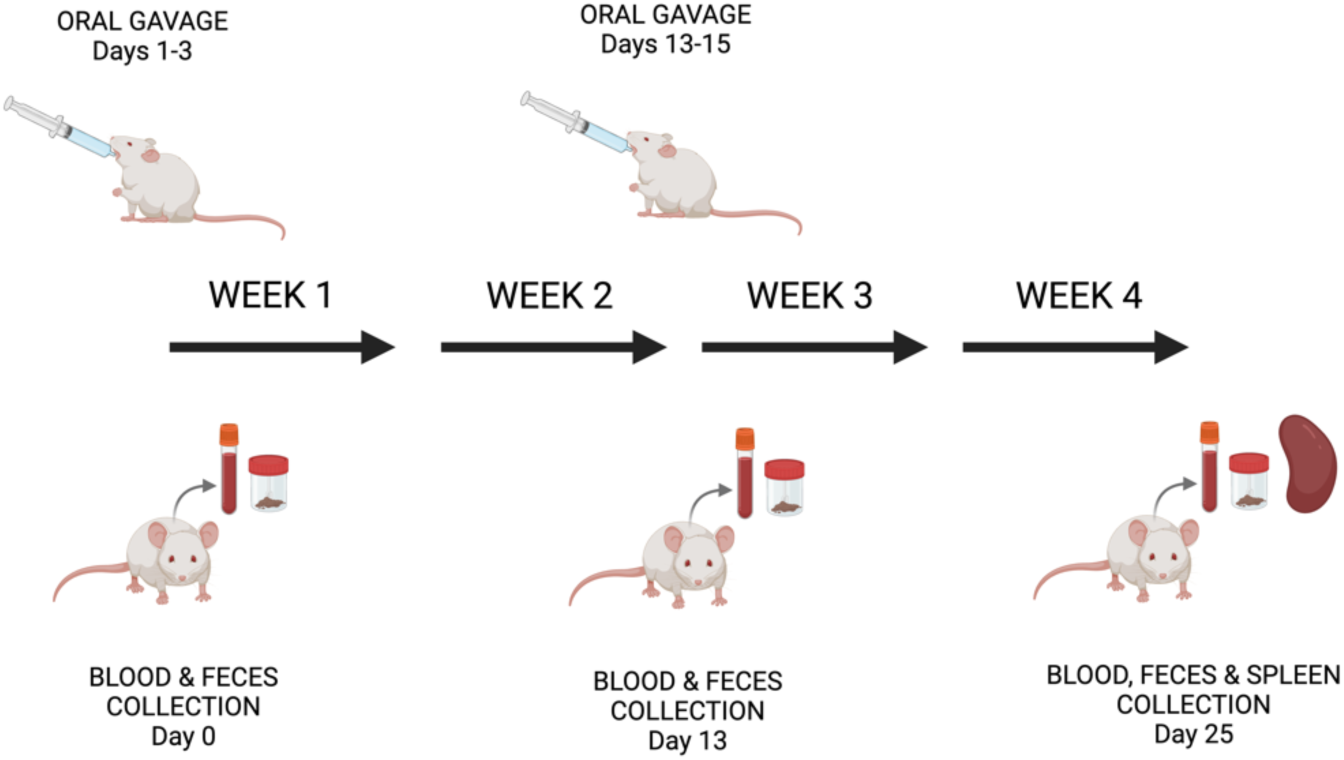
Oral Vaccination Schedule. On days 1-3, the four groups of mice received their designated strain of *S. boulardii* (2x10^9^ yeast cells/mouse) by oral gavage as a primary dose. Oral administration of the yeast was repeated on days 13-15 as a booster dose. Blood samples and feces were collected from all mice on days 0, 13, and 25. At the end of the fourth animal trial, spleens were also harvested on Day 25.

### Sample Collection and Immunological Analysis

Serum and fecal samples were collected on days 0, 13, and 25, corresponding to the administration of the primary dose on days 1-3 and the booster dose on days 13-15, respectively. Dried fecal pellets (100μg) were resuspended in 1mL of PBS-T (0.05% Tween) with 1μL of 10mM PMSF. The samples were vortexed thoroughly and incubated overnight at 4°C with constant shaking. They were centrifuged at 14,000 RPM for 10 minutes, and the final supernatants were used as fecal extracts. The titer of RBD-specific antibodies in serum and in fecal extracts was determined by ELISA.

### RBD-Specific IgG and IgA ELISA

For ELISAs, a Costar 96-well plate (Corning) was coated (200μL/well) with SARS-CoV- 2 Spike RBD-His Recombinant Protein (RayBiotech, USA) in carbonate coating buffer (pH 9.6) overnight at 4°C with gentle shaking. The concentration of RBD used was 2μg/mL. The plates were washed three times with PBS-T (0.05% Tween) and blocked (200 μL per well) with blocking buffer (PBS-T containing 1% BSA) at room temperature for 2 hours. Next, depending on the experiment, each was incubated either with 20x diluted sera in blocking buffer (100 μL) or fecal supernatants (100 μL) at 37°C for 1 hour. The ELISA plates were then washed three times with TBS-T (0.05% Tween), and, depending on the experiment, incubated either with biotinylated goat anti-mouse IgG (R&D Systems, USA) or HRP-conjugated goat anti-mouse IgA (Invitrogen, USA) at a 1:5000 dilution in TBS-T at 37°C for one hour. Again, they were washed 3x with TBS-T (0.05% Tween).

Finally, depending on the experiment, each well was incubated with either 100μL of alkaline phosphatase (AP)-conjugated streptavidin (R&D Systems, USA) at a 1:1000 dilution or 100μL of 3,3′,5,5′-tetramethylbenzidine (TMB) substrate at room temperature for one hour. Additionally, following an extra washing step (3 times with TBS-T), ELISAs for IgG titers were incubated with 100μL of pNPP phosphatase substrate (MP Biomedicals, USA) at room temperature for 30 minutes to develop the reaction. Measurement of absorbance was performed using a microplate reader (Bio-Tek, USA) at either 405nm (for IgG analysis) or 450nm (for IgA analysis). Note that for the third and fourth animal trials, we were able to complete the necessary measurements to express Ig values as endpoint titers. We had not done this for the first two trials.

### RBD-Specific IgG and IgA ELISA

During our fourth animal trial, splenocytes were isolated from vaccinated mice at the end of the experiment on day 25. Splenocytes (1× 10^6^ cells/well) were cultured in RPMI 1640 phenol red-free medium supplemented with 10% (v/v) FBS, 1 mM sodium pyruvate, 1 mM sodium glutamate, and a 1X antibiotic-antimycotic solution. Splenocytes were incubated for 72 hours with 2µg/mL SARS-CoV-2 Spike RBD-His Recombinant Protein (RayBiotech, USA). After the 72- hour challenge, culture spent media were collected to determine the interferon gamma (IFN-γ) and interleukin-4 (IL-4) secretions of the RBD-specific splenocytes using a mouse IFN-γ and mouse IL-4 ELISA kits (Solarbio, Inc., China). In brief, 100µL of spent medium and standard solution were pipetted into 96-well plates. The coated plates were incubated for one hour with 100µL of Biotinylated Mouse Anti-IFN-γ or Anti-IL-4 detection antibody, followed by a 45-minute incubation of 100µL Streptavidin-HRP at room temperature. An addition of 100 µL of TMB substrate was done and incubated for 30 minutes at room temperature in a dark room. The reactions were stopped by adding 50 µL of the stop solution, followed by an absorbance reading at 450 nm using a microplate reader. Each step was washed three times with a wash buffer.

### Statistical Analysis

Statistical analysis of the total IgG and IgA levels was performed using Prism version 10.0.3 (GraphPad Software, USA). Differences between the experimental and control groups were analyzed using an unpaired Student’s t-test. P-values of <0.05 were considered statistically significant.

## Results

To create our candidate yeast oral vaccine, we chose to utilize the probiotic yeast *Saccharomyces boulardii*, which has been approved for human consumption worldwide (21), as a vaccine delivery system. Interestingly, data suggest that although *S. boulardii* induces a systemic humoral immune response, this response is small in magnitude and not directed against *S. boulardii* itself (22). In principle, this suggests that the probiotic yeast could serve as a relatively immunologically benign vaccine platform for the repeated administration of various target antigens.

To express our SARS-CoV-2 target antigen, we designed a construct with an integrative yeast (YIp) plasmid that would express the Receptor Binding Domain (RBD) of the Spike protein of SARS-CoV-2 from the strong constitutive TEF1 yeast promoter. The RBD is the most immunogenic epitope of the virus (23), and the TEF1 yeast promoter is known to function robustly in *S. boulardii* (19, 24). To target the gut-associated lymphoid tissue (GALT), we fused the 125 amino acid C-terminal fragment of the *Clostridium perfringens* enterotoxin (CpE) to the C-terminus of the RBD domain. This C-terminal fragment of the CpE targets proteins of the gap junctions of cells found in the GALT (26). It has also been used to induce an immune response against ovalbumin in the gut of mice (19). These studies suggest that the CpE system could be used to facilitate the interaction between the SARS-CoV-2 antigen and the gut’s immune cells. Next, to force the yeast cells to secrete our target antigen, we fused the 89-amino-acid yeast α- factor mating secretion signal to the N-terminus of the 217-amino-acid RBD domain (25). This yeast mating pheromone signal is cleaved from the protein just before secretion from the cell. In toto, we created a construct that could be integrated into yeast at the URA3 locus in order to express and secrete a target antigen composed of an RBD-CpE fusion protein.

Next, several *Saccharomyces boulardii* strains transformed with linearized versions of our construct were cultured for 48 hours and screened for secretion of the SARS-CoV-2 RBD antigen using standard Western blot analysis. As shown in Figure 2, we identified two independent yeast vaccine strains that express and secrete the Wuhan RBD-CpE fusion protein into the culture medium, which we named W2 and W5. The upper band in our Western blot appears to correspond to the RBD-CpE fusion protein with the 89-amino-acid yeast α-factor mating secretion signal, which is expected to have a size of 41kD. In contrast, the lower band represents the fusion protein without the secretion signal, which is cleaved just before secretion from the yeast cell (26). Both of these yeast vaccine strains were used to immunize BALB/c mice to see if our candidate vaccine could elicit a detectable immune response.

**Figure 2.**
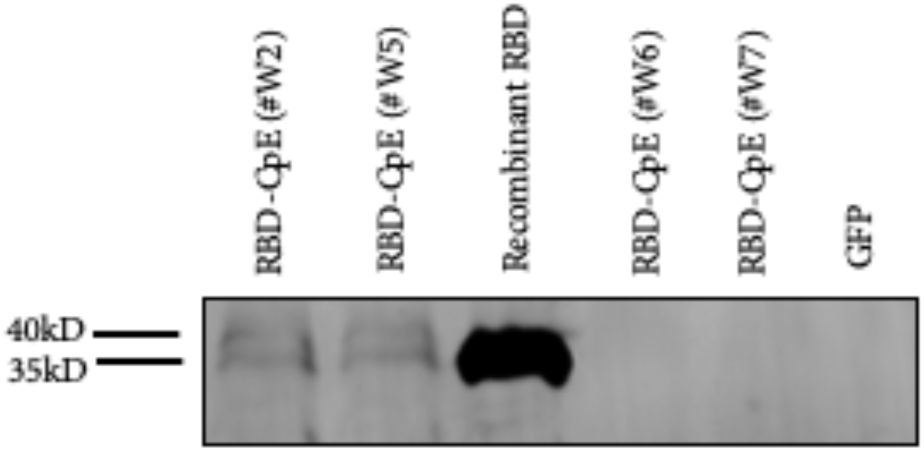
Recombinant Strains of Saccharomyces boulardii Secrete the RBD-CpE Fusion Protein. Western blot analysis of four recombinant S. boulardii strains reveals that two strains (W2 and W5) of the four tested candidates secrete the RBD-CpE fusion protein into the culture media.

To determine whether or not the oral administration of our candidate yeast oral vaccine could elicit detectable IgG and IgA responses in mice, we completed four independent animal trials. The first two trials included cohorts of BALB/c mice inoculated with *Saccharomyces boulardii* alone, with *S. boulardii* expressing GFP, or with one of two candidate vaccine *S. boulardii* strains (W2 or W5) expressing and secreting RBD-CpE into the culture media. Given the results of the first two animal trials, which suggested that both recombinant yeasts were comparable and that both negative controls were comparable, we decided to limit our third and fourth animal trials to the W5-RBD vaccine mice alone, with a wild-type M2 yeast with no insert alone as a negative control. Notably, none of our mice showed any signs that our yeast or yeast vaccine was detrimental to their well-being. In fact, as shown in Figure 3, the weights of the mice were comparable throughout the duration of the third trial.

**Figure 3.**
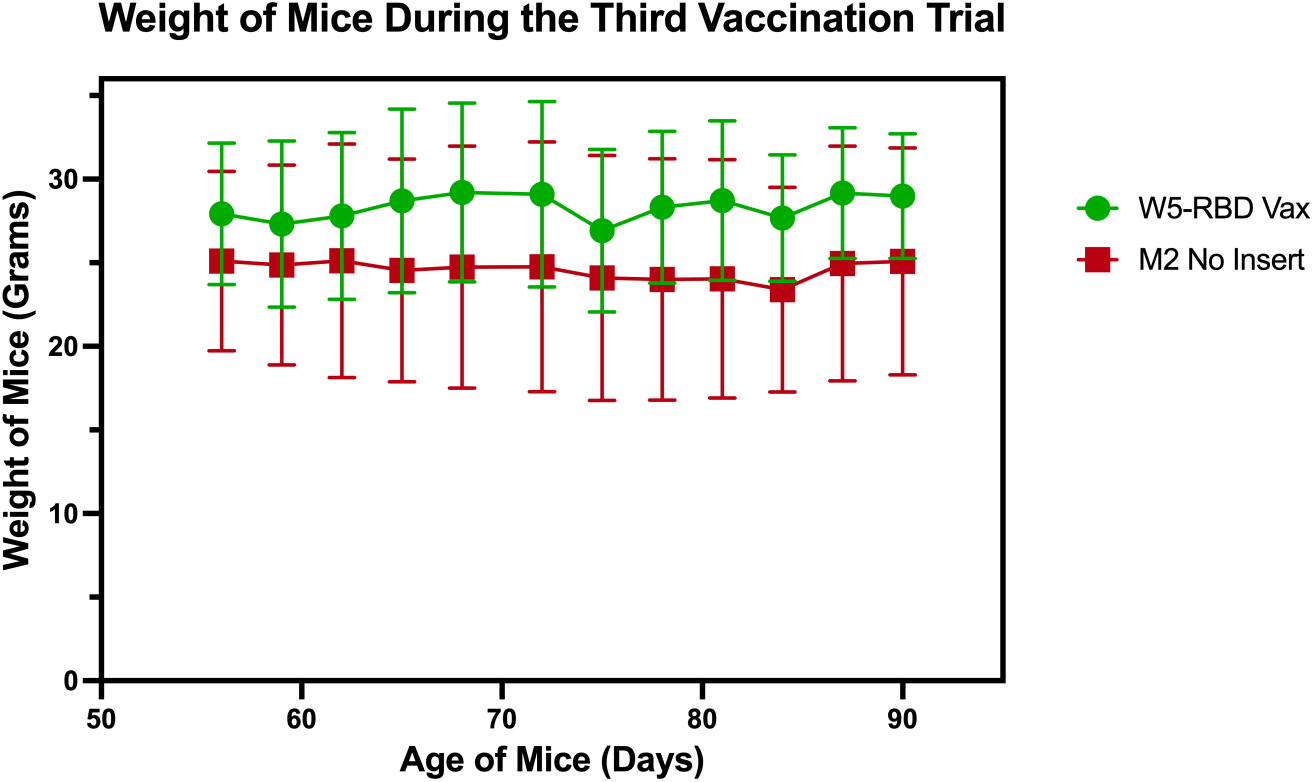
BALB/c Mice Fed Either *Saccharomyces boulardii* Alone or *S. boulardii* Expressing the RBD-CpE Fusion Have Similar Weights Throughout the Course of the Month-long Experiment. Mice fed either *Saccharomyces boulardii* Alone or *S. boulardii* Expressing the RBD-CpE were weighed every other day during the course of the third vaccination trial. The figure shows mean weights of the two cohorts of mice along with their standard deviations. The differences were not statistically significant at all times tested.

As shown in Figure 4, four independent animal trials reveal that recombinant strains of *Saccharomyces boulardii* secreting the RBD-CpE fusion protein can elicit a detectable IgG response against the SARS-CoV-2 RBD. IgG serum levels were significantly elevated (p<0.005) in the cohorts of vaccinated mice two weeks and four weeks after the primary dose, as compared to those mice that received yeast alone or yeast with GFP.

**Figure 4.**
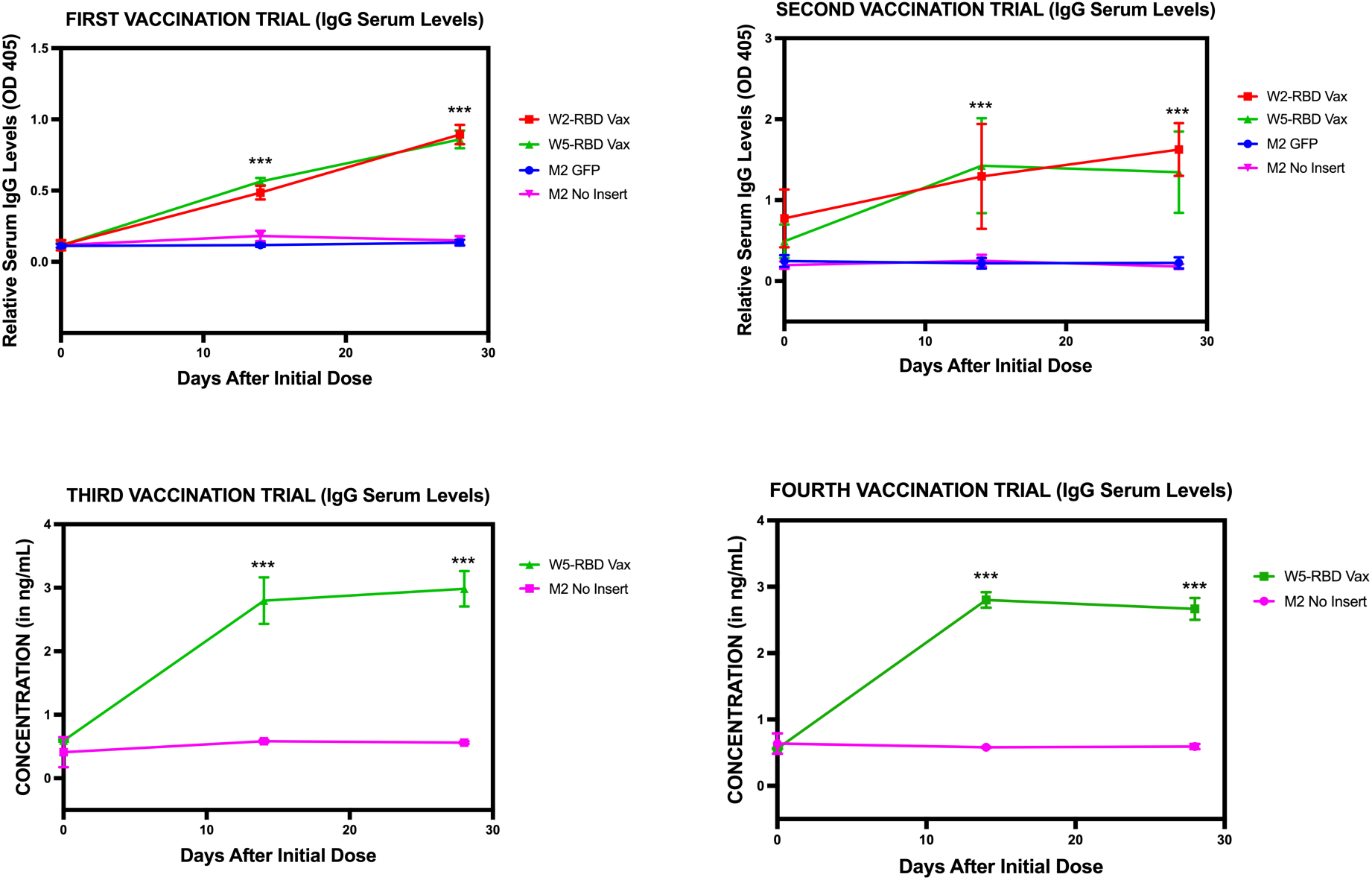
Recombinant Strains of *Saccharomyces boulardii* Secreting the RBD-CpE Fusion Protein Can Elicit a Detectable IgG Antibody Response against SARS-CoV-2 RBD. Serum from mice in four independent trials was analyzed using an ELISA to determine if it contained IgG antibodies against the SARS-CoV-2 RBD. The first two trials included cohorts of BALB/c mice inoculated with *Saccharomyces boulardii* alone, with *S. boulardii* expressing GFP, or with one of two candidate vaccine *S. boulardii* strains (W2 or W5) expressing and secreting RBD-CpE into the culture media. Given the results of these initial two trials, our third and fourth animal trials were limited to W5-RBD vaccine mice alone, with a wild-type M2 yeast with no insert alone as a negative control.

Next, as shown in Figure 5, the second, third, and fourth animal trials also demonstrated that recombinant strains of *Saccharomyces boulardii* secreting the RBD-CpE fusion protein can elicit a detectable IgA response against the SARS-CoV-2 RBD. (We encountered technical difficulties processing the feces we had collected from the mice during the first animal trial.) Our data revealed that IgA levels were significantly elevated (p<0.005) in the fecal extracts of cohorts of vaccinated mice two weeks and four weeks after the primary dose, as compared to the extracts of those mice who had received yeast alone or yeast with GFP. IgA levels are associated with mucosal immunity, which we would expect to be triggered by an oral vaccine.

**Figure 5.**
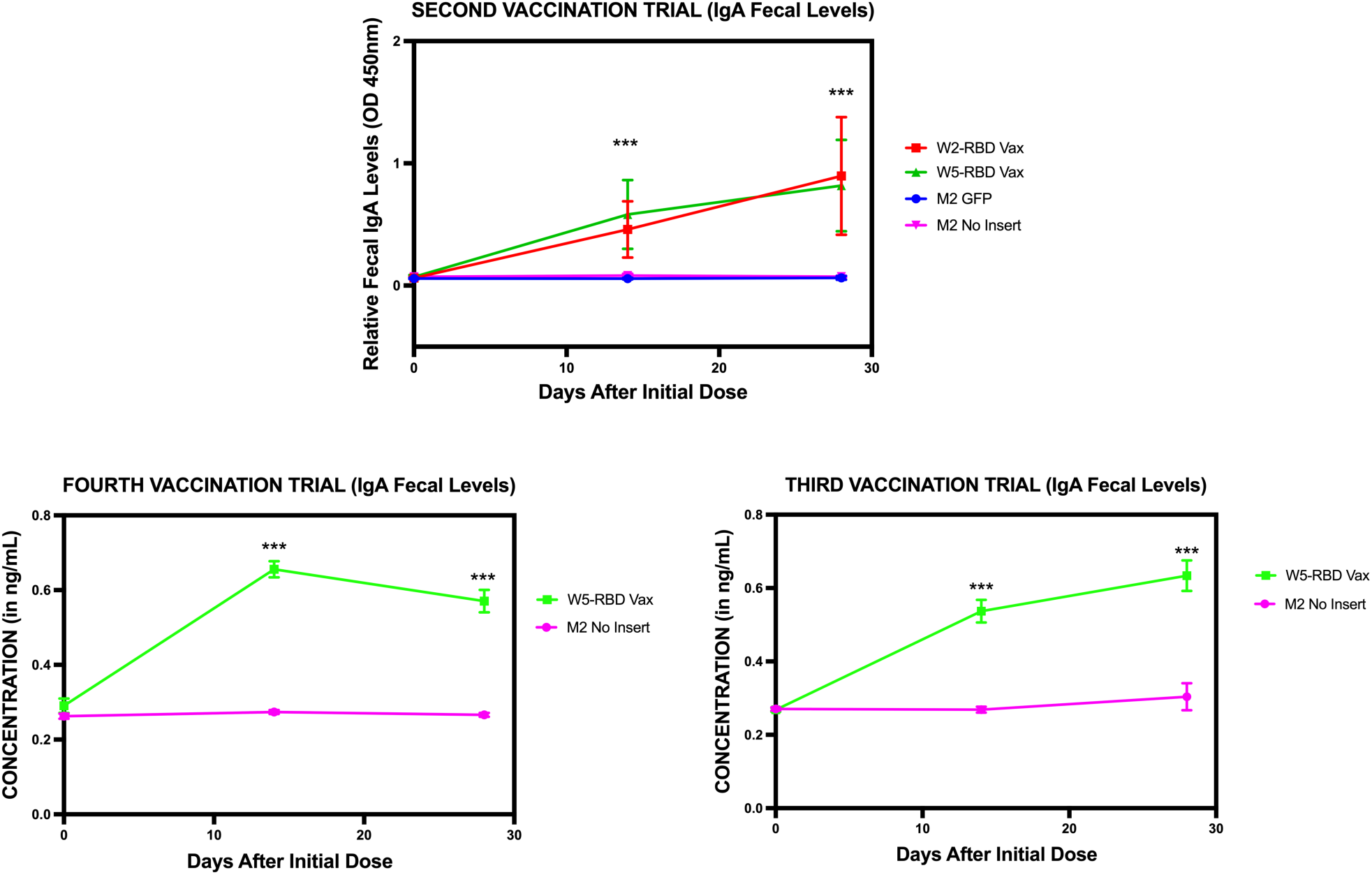
Recombinant Strains of Saccharomyces boulardii Secreting the RBD-CpE Fusion Protein Can Elicit a Detectable IgA Antibody Response against SARS-CoV-2 RBD. Fecal extracts of mice from three vaccination trials were analyzed with an ELISA to determine if they contained IgA antibodies against the SARS-CoV-2 RBD. The second trial included cohorts of mice that were inoculated with Saccharomyces boulardii alone, with S. boulardii expressing GFP, or with one of two candidate vaccine S. boulardii strains expressing and secreting RBD-CpE into the culture media. Given the results of this initial trial, our third and fourth animal trials were limited to W5-RBD vaccine mice alone, with a wild-type M2 yeast with no insert alone as a negative control.

Finally, as shown in Figure 6, we were able to isolate splenocytes from our mice after the fourth animal trial and to challenge them with recombinant SARS-CoV-2 RBD to determine if they would respond to the antigen. (Difficulties in procuring reagents because of the pandemic precluded us from performing these experiments sooner.) Splenocytes obtained from vaccinated mice secreted increased levels of IFN-γ but not of IL-4 when exposed to SARS-CoV-2 RBD. This suggests that our yeast vaccine candidate is capable of activating the cell-mediated arm of the immune system. Significantly, the absence of an IL-4 response indicates that the vaccine response is Th1-biased, favoring cell-mediated immunity over an antibody-dominated response.

**Figure 6.**
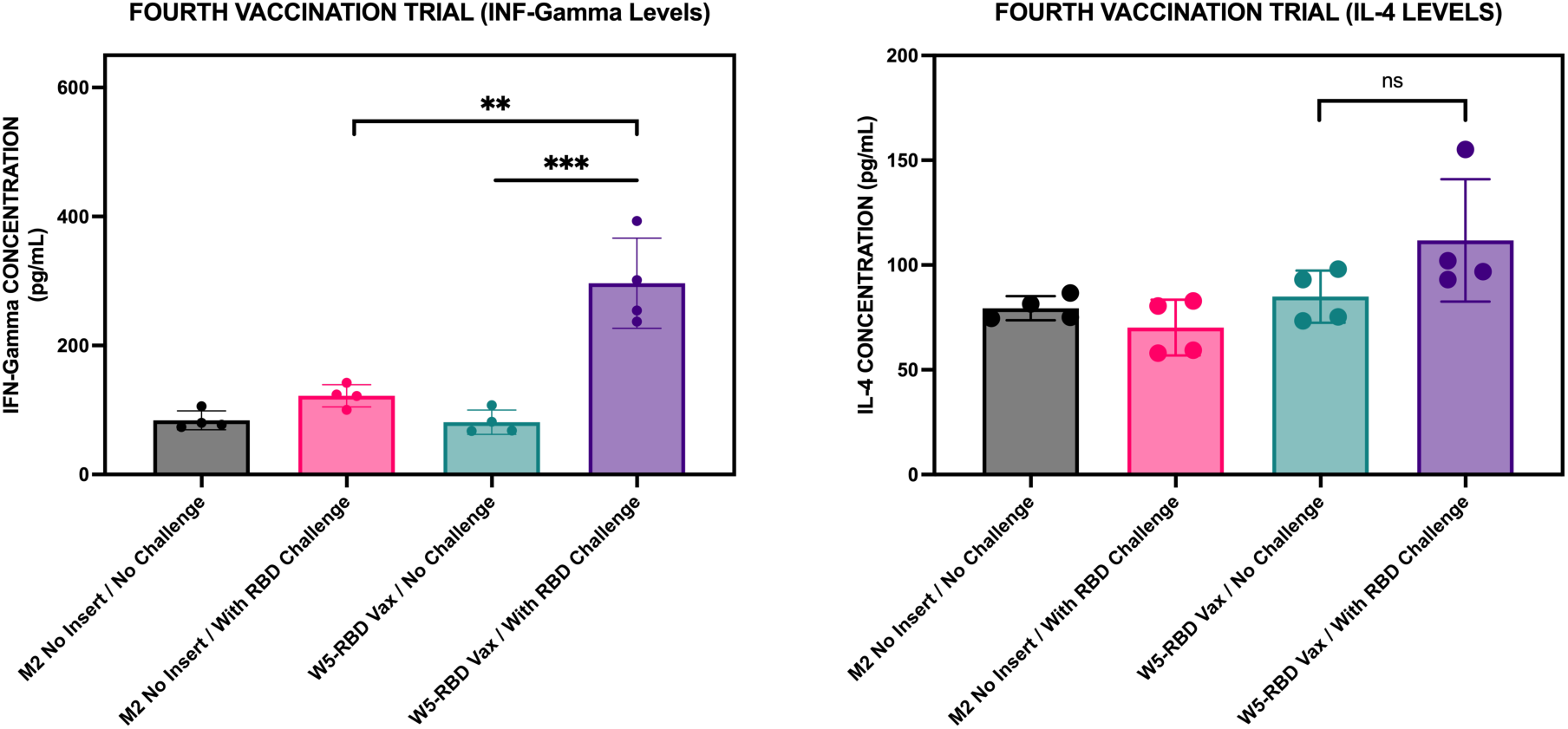
Recombinant Strains of Saccharomyces boulardii Secreting the RBD-CpE Fusion Protein Can Elicit a IFN-γ but not IL-4 Response against SARS-CoV-2 RBD. Splenocytes of mice from our fourth animal trial were isolated and challenged with recombinant RBD. Cytokine responses in the spent media were measured with an ELISA. The cohorts included mice that were inoculated with Saccharomyces boulardii alone or with a candidate S. boulardii vaccine strain expressing and secreting RBD-CpE

In summary, our data suggest that probiotic yeast expressing our secreted SARS-CoV-2 RBD-CpE yeast candidate oral vaccine can induce a detectable antibody response, including both IgG and IgA, as well as a cell-mediated response that is Th1-biased in BALB/c mice. As such, our experiments suggest that *Saccharomyces boulardii* remains a promising and novel vaccine delivery platform not only for COVID-19 but also for other infectious diseases.

## Discussion

Over the past decade, various research groups have conducted studies to investigate the potential use of yeast cells as a vaccine delivery platform for the oral administration of target antigens (8). There are at least three inherent advantages that yeast delivery platforms have over other vesicular, viral, or bacterial systems. First, numerous clinical trials have demonstrated that Saccharomyces cerevisiae and its conspecific probiotic relative, Saccharomyces boulardii, are safe for human consumption (21, 27, 28). Second, these yeasts survive transit through the harsh environment of the mammalian gut. When S. boulardii was given to healthy volunteers as a probiotic at therapeutic doses (1–2 × 10^10^/d), colonic levels were 2 × 10^8^/g stool (29). One possible reason for this is that S. boulardii cells are resistant to low pH and bile acid (16, 30–32). Finally, these yeasts are known to stimulate the immune system, making them natural adjuvants (33, 34). Together, these characteristics suggest that yeast cells are prime candidates for an oral vaccine delivery system.

Despite the declaration of the WHO that the COVID-19 pandemic has ended, vaccine development remains an important task for future outbreaks, especially in lower- and middle-income countries like the Philippines, where vaccine scarcity is commonplace. In this paper, we report the successful engineering of recombinant strains of Saccharomyces boulardii to express and secrete the Receptor Binding Domain (RBD) of the Spike protein from the original Wuhan-Hu-1 strain of the SARS-CoV-2 virus, fused to the C-terminal fragment of Clostridium perfringens enterotoxin (CpE). Animal trials suggest that our candidate vaccine can elicit a detectable adaptive immune response in BALB/c mice. This response is characterized by increased production of IgG and IgA, as well as enhanced expression of IFN-γ from splenocytes isolated from vaccinated mice.

Our vaccine candidate is not the first yeast COVID-19 vaccine described. While we were conducting our experiments during the pandemic lockdown, two separate scientific teams reported that they had developed candidate yeast oral vaccines for COVID-19 using Saccharomyces cerevisiae (20, 35). These candidate COVID-19 vaccines also used the receptor-binding domain (RBD) of the SARS-CoV-2 spike protein as a target antigen. In one case, the RBD domain was also fused to the adjacent fusion peptide (FP) domain (35).

However, unlike our vaccine, which was designed to secrete the target antigen, these other COVID-19 yeast oral vaccines used surface display technology to coat the surface of their yeast cells with the target antigen. Cell-surface display directs the expression of target peptides or proteins to the cell surface of a diverse range of cell types through the connection of a protein of interest fused to an anchor protein (36). Nonetheless, like our secreted vaccine candidate, both candidate vaccines using cell-surface display were able to elicit detectable IgG and IgA responses against the SARS-CoV-2 RBD domain.

One research group also demonstrated that their cell-surface-displayed candidate vaccine was capable of triggering detectable cellular immune responses, as evidenced by T-cell proliferation accompanied by increased IFN-γ and IL-4 expression upon exposure to the antigen (20). In contrast, our secreted candidate vaccine was able to induce an IFN-γ but not an IL-4 response. It is not clear why this is the case. Interestingly, studies in BALB/c mice, which are inherently Th2-biased, have shown that specific IM vaccine formulations can increase IFN-γ production with no significant change in IL-4 levels (46, 47). The authors of both studies speculate that the Th1-biased response could be explained by CpG motifs in their plasmid backbone and the use of intracellular Toxoplasma antigens, both of which drive strong Th1 polarization and suppress Th2 cytokine induction.

In light of these differences, it will be essential to conduct side-by-side experiments with identical dosages and vaccine schedules to determine whether one oral yeast vaccine design, either secreted or cell-surface-displayed, is more effective than the other. Moreover, our COVID-19 vaccine candidate includes a CpE targeting signal (13–15). We included this element in our vaccine design to enhance the interaction between the RBD and the gut-associated lymphoid tissues (GALT). Since the RBD of SARS-CoV-2 can bind to ACE2 in the GALT (37), it should be interesting to see if deleting the CpE mitigates the humoral and cell-mediated responses that we have detected. If CpE is crucial for immunogenicity, a shorter fragment of the CpE could also be tested for function (38).

Notably, elevated immune responses were observed among Saccharomyces cerevisiae COVID-19 vaccine candidates, even though two distinct vaccination schedules were employed. For one study, mice were orally administered the yeast vaccine on days 1 and 2 as a primary dose and then again on days 14 and 15 as a booster (20). This schedule approximates the schedule utilized in our study. In contrast, the second study reported that their mice received oral vaccines on day 1 for priming immunization and then again on days 5, 10, and 21 for boosting immunization (35). Despite the distinct vaccination schedules used, however, all three vaccine candidates, including ours, were able to trigger a detectable humoral response. For two of them, including ours, a detectable cell-mediated response against the SARS-CoV-2 RBD was also observed. Together, these three studies, ours included, suggest that the precise schedule for vaccinating the mice may not be critical for vaccine efficacy.

One of the primary drawbacks of an oral vaccine is that higher doses of antigen or multiple doses are required to compensate for the loss of antigen in the harsh gut environment (39). To address this challenge, we plan to optimize our vaccination schedule in future experiments, particularly since data suggest that altering the interval between priming and boosting can modify the host immune response profile (40). Moreover, we also intend to test whether adjuvants can enhance the efficacy of our vaccine candidate. Once again, due to the lower bioavailability of antigens in oral vaccine formulations, adjuvants are often required to make an oral vaccine effective (41).

Another challenge associated with oral vaccines is that their efficacy can be mitigated by the gut microbiota (39). Although the gut microbiome can influence vaccine-induced immunity in humans, the causal mechanisms underlying this phenomenon remain unclear (42, 43). One study conducted in Bangladesh has revealed that the abundance of Bifidobacterium during early infancy is correlated with the immune response to the oral polio vaccine, among other factors, and that this correlation persists until the child is two years old (44). Our yeast oral vaccine candidate utilizes the probiotic yeast Saccharomyces boulardii as a delivery platform. Studies suggest that probiotics can enhance the oral immune response to rotavirus in a porcine model (45). It will be interesting to interrogate the impact of this yeast probiotic on the efficacy of our oral vaccine and to determine if the delivery platform itself can act as an adjuvant.

Finally, like our study, neither of the published studies describing the Saccharomyces cerevisiae COVID-19 vaccine candidates reported the levels of neutralizing antibodies against SARS-CoV-2, and neither COVID-19 vaccine was the subject of a direct challenge animal trial. Both of these will have to be assessed for all three vaccine candidates, including ours, if any of them are to proceed to human clinical trials.

## Ethics Statement

This study was approved by the Institutional Animal Care and Use Committee (IACUC) of the University of Santo Tomas, Manila, Philippines, on September 28, 2022, with protocol number RC2021-030723.

## Author Contributions

Conceptualization: NA; experimental analysis: JDAR, CB, TB, NK, GM, LCM, MKD, JRC, DAEC, JPA, MGR, and NA; data analysis: JDAR, CB, TB, NK, and NA; writing and revising of manuscript: JDAR, TB, NK, GM, MKD, DAEC, and NA; funding acquisition: NA. All authors have read and agreed to the published version of the manuscript. JDAR, CB, TB, and NK contributed equally to this work.

## Acknowledgments

We thank Marc Martin Sagolili for technical assistance. We also thank Milo B. Fasken and Tracey J. Lamb (Emory University, USA) for their generous gift of the M2 strain of Saccharomyces boulardii, Vahid Khalaj (Pasteur Institute of Iran, Iran) for his expert advice at the outset of this study, and our anonymous peer reviewers for their comments. The images of the oral vaccination schedule and the graphical abstract were created with Biorender.com.

## Current Affiliations

CB is currently at the Dominican House of Studies in Washington, DC 20017 USA; TB is presently at the Department of Systems Pharmacology and Translational Therapeutics, Perelman School of Medicine at the University of Pennsylvania, Philadelphia, PA 19104 USA; NK is currently at the Sellers Laboratory at the Broad Institute of MIT and Harvard, Cambridge, MA 02142 USA; and JPA is at the Division of Biology and Biomedical Sciences at Washington University, St. Louis, MO 63130 USA.

